# The Gordian Knot Enhances Ubiquitin Binding in UCH-L1

**DOI:** 10.64898/2026.08.29.747984

**Authors:** Sara G. F. Ferreira, Patrícia F. N. Faísca, Miguel Machuqueiro

## Abstract

UCH-L1 is a monomeric deubiquitinating enzyme whose native structure embeds a shallow 5_2_ knot near the N-terminus, placing the knotted topology in direct proximity to both the substrate-binding pocket and the catalytic site. While our previous work ^1^ established that N-terminal integrity is critical for catalytic activity, the energetic and structural consequences of removing the knot without altering the amino acid sequence remained unresolved. Here, we combine steered molecular dynamics with umbrella sampling to generate full-length, sequence-identical variants of UCH-L1 with weakened or disrupted topology and reconstruct the free-energy profile along a constrained unthreading pathway. The potential of mean force places the sampled unknotted conformations approximately 5–7 kcal/mol above the native knotted minimum, indicating a thermodynamic preference for the native topology along this pathway. Long unbiased MD simulations of fully unknotted variants in both apo and holo states show that knot removal predominantly increases local N-terminal flexibility without significantly destabilizing the global fold or disrupting the catalytic-site geometry. MM-PBSA calculations further predict less favorable ubiquitin-binding energetics upon unknotting (*∼*-62 vs *∼*-76 kcal/mol), suggesting that topology contributes to substrate affinity. Together, these results suggest that the 5_2_ knot in UCH-L1 is not a passive topological feature but a functional element that constrains unfolding dynamics and contributes to substrate binding efficiency.

## Introduction

Knotted proteins are proteins whose native structure embeds a physical (i.e., open) knot. Advances in knot-detection algorithms and loop-closure procedures ^2,3^ have enabled systematic analysis of protein structures, leading to the discovery of several knot types.^4,5^ Like topological knots (which are closed curves in space), protein knots can also be classified based on the minimal number of crossings of a planar projection of the knot. Knotted proteins offer unique opportunities to study the interplay among topology, folding, and biological function.^6–10^

An interesting example of a knotted protein is ubiquitin C-terminal hydrolase L1 (UCH-L1). UCH-L1 is a single-domain deubiquitinating enzyme with 223 amino acids that features a Gordian (5_2_) knot in its native structure. The knot spans the substrate-binding pocket and the catalytic site and is shallow, with the knot core located only 4 residues from the N-terminus. UCH-L1 is highly expressed in neuronal tissues, accounting for 1 to 5% of total neuronal protein content,^11^ and in several forms of cancer.^11,12^ Axonal integrity strictly depends on UCH-L1, and mutations in UCH-L1 have been linked to neurodegenerative disorders, such as Alzheimer’s and Parkinson’s disease. ^13–15^ The physiological and pathological importance of UCH-L1, along with its native knotted structure, has driven a body of experimental and computational work to understand its folding and function.

Bulk experiments revealed that UCH-L1 populates two parallel folding pathways, each featuring a metastable unknotted intermediate.^16,17^ Similar observations were reported in molecular simulations using a structure-based model.^18^ A far more complex picture emerged from single-molecule experiments. Indeed, optical tweezers revealed that the folding landscape of UCH-L1 features many on- and off-pathway intermediate states during unfolding and refolding, and showed that the formation of a 3_1_ or 5_2_ knot significantly slows folding of UCH-L1.^19^ These observations are consistent with the fact that knotting, which requires threading the shortest knot terminus through a transient loop formed by the remainder of the chain (as reviewed in^20^), is a rate-limiting step in the folding of tangled proteins. Interestingly, the single-molecule experiments also revealed that, at low to moderate forces, the 5_2_-knotted region in denatured conformations of UCH-L1 is considerably large, spanning about 40 residues, thereby delaying proteasomal processing due to steric clashes within the narrow proteasome translocation channel. ^19^

A subsequent study that employed the bacterial AAA^+^ protease system ClpXP to evaluate how UCH-L1, amongst other knotted deubiquitinases, resists mechanical unfolding and proteolysis reported a remarkably long lifetime under ClpXP-mediated degradation (1039 min) for UCH-L1.^21^ This observation strengthened the view that the knot adds mechanical stability and protection from unfolding and degradation.

Recently, we combined molecular dynamics simulations with *in vitro* experiments to explore the role of the UCH-L1 knot in catalytic activity. ^1^ In doing so, we serendipitously found that truncating only 2 residues from the N-terminus significantly affects the catalytic activity of UCH-L1 without altering its secondary structure or topological state. As expected, truncating 5 residues unties the protein. Additionally, it introduces significant changes to the secondary structure and completely abolishes catalytic activity. While these results do not allow us to claim that the knot *per se* plays a direct role in catalytic activity, they strongly suggest that it may contribute indirectly by stabilizing the overall structure and thereby enabling the correct alignment of the catalytic triad. However, because unknotting was achieved by N-terminal truncation, the effects of changing topology could not be separated from those caused by altering the amino acid sequence. Therefore, the energetic cost and structural consequences of removing the knot while preserving the full-length sequence remained unresolved.

Here, we employ atomistic molecular dynamics simulations coupled with an umbrella sampling scheme to generate full-length, sequence-identical variants of UCH-L1 in which the knot is disrupted or weakened. This strategy allows us to distinguish the consequences of changing topology from those associated with N-terminal truncation. We characterize the free-energy profile along the sampled unthreading pathway and investigate how knot removal affects the global fold, local N-terminal dynamics, catalytic-site geometry, and calculated ubiquitin-binding energetics.

## Materials and Methods

### System Preparation

In this work, we considered the apo state of UCH-L1 (PDB ID: 2ETL^22^) and its holo complex with ubiquitin (PDB ID: 3KW5^23^). The terminal residue of ubiquitin (Gly76), which was unresolved in the crystal structure, was rebuilt using the Builder tool in PyMOL.^24^ All simulations preserved the full 223-residue sequence of UCH-L1.

Before solvation, protonation states for all titratable residues were assigned based on p*K*_a_ estimates obtained from the PypKa server.^25,26^ No unusual protonation states were predicted at pH 7.0; therefore, all histidines were kept neutral. Systems were solvated in a periodic dodecahedral box using the SPC water model. ^27^ For the apo system, 6500 water molecules and 8 Na^+^ ions were added to achieve charge neutrality, whereas the holo system required approximately 14000 water molecules and 8 Na^+^ ions. Steered-MD-derived systems were solvated analogously, with minor adjustments to the number of water molecules and neutralizing ions to account for the altered spatial dimensions of the pulled conformations. Since both apo and holo starting structures had already undergone complete energy minimization and equilibration in our previous MD study,^1^ no additional steps were required before initiating the steered MD simulations. The structures used here correspond directly to equilibrated endpoints of those validated trajectories, ensuring methodological continuity across studies.

### Steered Molecular Dynamics

To investigate the structural and energetic constraints imposed by UCH-L1’s knotted topology, we designed a steered molecular dynamics (steered-MD) protocol to progressively untie the knot by extracting the N-terminus of UCH-L1 from its knotted core. This approach also enabled the generation of topologically modified UCH-L1 variants that retained their full-length sequence, which were essential for subsequent umbrella sampling calculations.

Pulling was applied to the C*_α_* atom of a selected N-terminal residue, while all other protein atoms were harmonically restrained except for residues 1–10 of UCH-L1, which were left entirely unrestrained. In the holo simulations, all ubiquitin atoms were likewise restrained to preserve the structural integrity of the bound complex during pulling.

The pulling coordinate was defined as the distance between the center of mass of the gate residues (88, 140, 144, 147, 153, 155, and 157) and the selected N-terminal residue (Figure 1). Although the selected collective variable cannot capture all microscopic degrees of freedom involved in knot disruption, it directly monitors the passage of the N-terminus through the gate. It therefore represents the dominant geometric event required for unknotting. Pulling was applied along the distance between these two groups using a constant-velocity protocol with a pulling rate of 0.0001 nm ps*^−^*^1^ (0.1 nm ns*^−^*^1^).^28^ Apo simulations used a force constant of 1000 kJ mol*^−^*^1^ nm*^−^*^2^, whereas holo simulations required a higher force constant of 5000 kJ mol*^−^*^1^ nm*^−^*^2^ due to steric hindrance from the C-terminus of bound ubiquitin near the gate region.

**Figure 1:**
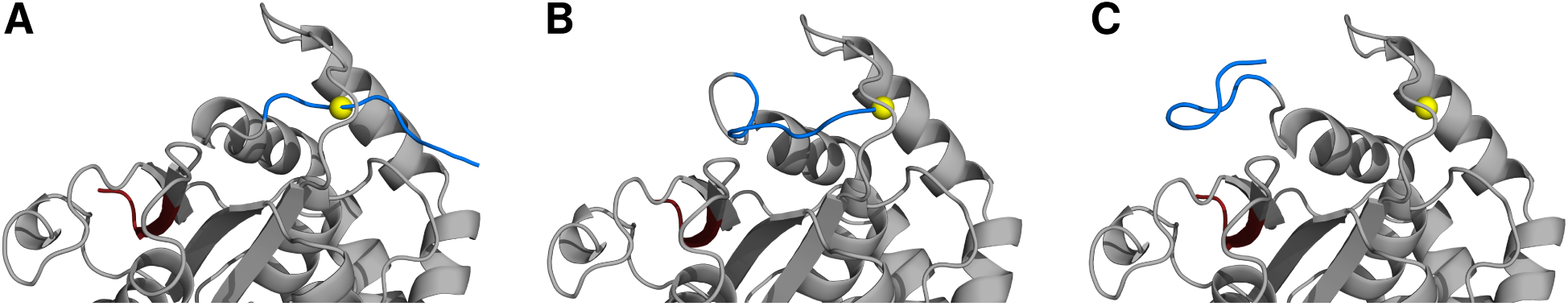
Illustration of the UCH-L1 conformational transition obtained in the steered MD protocol. The starting conformation (A) corresponds to the X-ray structure (PDB ID: 2ETL^22^) where the protein is fully knotted. At half-transition (B), the N-terminus is located exactly under the gate loop. At the end of the steered MD protocol (C), the N-terminus region is fully unknotted. The protein is represented as a gray cartoon. The N-terminal region (7 initial residues) is colored in blue, the C-terminal region (7 final residues) is colored in red, and the center of mass of the gate residues (88, 140, 144, 147, 153, 155, and 157) is depicted as a yellow sphere.

Since the N-terminus needed to be pulled sequentially through the gate, the steered-MD protocol was performed in five consecutive 3-ns segments, yielding a cumulative pulling time of 15 ns. After each 3-ns segment, the identity of the pulled residue was updated to the next residue along the N-terminus: Glu7 (2*×*) *→* Met6 *→* Pro5 *→* Lys4, and the simulation was continued from the final coordinates and velocities of the preceding segment. Each segment generated a displacement of *∼*3 Å, enabling smooth extraction of the N-terminus and minimizing abrupt structural distortions. Intermediate conformations harvested from these steered-MD trajectories were subsequently used to initialize the umbrella sampling windows.

### Umbrella Sampling

Umbrella sampling (US) calculations were performed only for the apo state of UCH-L1. This choice was motivated by the fact that, in the holo complex, the C-terminal tail of ubiquitin partially occupies the gate region, sterically interfering with the reaction coordinate and preventing consistent sampling of both topological branches.

Intermediate structures extracted from the steered-MD trajectories were used to construct the reaction coordinate, defined as the distance between the C*_α_* atom of residue 1 (Met1) and the center of mass of the gate residues. Window 00 corresponded to the configuration in which the N-terminus was aligned with the gate (Figure 1B), whereas windows *−*16 (Figure 1C) to +16 (Figure 1A) sampled progressively displaced conformations up to 16 Å on either side of the central position, using 2 Å spacing between windows.

Because the reaction coordinate depends only on distance, it does not encode directional information. Therefore, to distinguish between the two pathways passing through the gate, we used PyMOL to construct a reference plane that intersects the gate’s center of mass. For each configuration, the position of residue 1 relative to this plane was used to classify the structure as belonging to the positive (+) or negative (-) branch of the reaction coordinate. The sign assignment was performed after the US simulations, before the WHAM calculation. Frames in which residue 1 lay very close to the plane could not be reliably assigned to either branch, as thermal fluctuations caused rapid side-switching in this region. Therefore, during the selection of starting structures for umbrella windows, we deliberately avoided configurations in which residue 1 was located within approximately 2 Å of the reference plane. This ensured that each umbrella window was seeded with a structure that unambiguously belonged to a single topological branch, preventing mixing of distinct pathways during the Weighted Histogram Analysis Method (WHAM) analysis.

Each umbrella window was simulated using five independent 100-ns replicates. Window 00 was simulated twice, once for the knotted and once for the unknotted configuration, yielding a total of ten replicates for the central window. In total, this resulted in 90 umbrella-sampling simulations. A harmonic umbrella force constant of 1000 kJ mol*^−^*^1^ nm*^−^*^2^ was applied in all windows.

### Unbiased Molecular Dynamics Simulations

To evaluate the structural stability of fully unknotted conformations, long unbiased MD simulations were performed for both apo and holo systems. For the apo state, 10 unknotted starting structures were selected from the umbrella window *−*8 (corresponding to an 8 Å unknotted displacement from the gate). These ten structures comprised two frames from each of the five US replicates and were simulated for 500 ns each.

For the holo state, a single fully unknotted conformation obtained from the steered-MD protocol was used as the starting structure for ten independent 500-ns replicates. Because steered MD continuously applies external forces, the extracted unknotted conformation has not previously been allowed to relax under unbiased equilibrium conditions. Therefore, a short equilibration was required before initiating the production runs. The system underwent a brief preparation stage comprising two steepest-descent minimization cycles, followed by restrained NVT and NPT equilibration (100 ps each). Harmonic restraints (1000 kJ mol*^−^*^1^ nm*^−^*^2^) were applied to all protein heavy atoms during equilibration and removed before the start of the unrestrained simulations.^29^ These steps ensured that the holo unknotted starting state was fully relaxed, enabling a reliable assessment of the stability, structural evolution, and topological integrity of unknotted full-length UCH-L1 in both binding states.

### Simulation Parameters

All simulations were carried out using the GROMACS 2021.2 package^30^ with the GROMOS 54A7 force field^31^ and the SPC water model.^27^ Electrostatic interactions were treated using the Particle-Mesh Ewald (PME) method,^32^ with a real-space cutoff of 1.4 nm and a Fourier grid spacing of 0.12 nm. Lennard-Jones interactions were truncated at 1.4 nm. All covalent bonds were constrained using the LINCS algorithm with lincs_order = 8.^33^ All simulations were performed with a 2 fs timestep.

Temperature was maintained at 310 K using the v-rescale thermostat^34^ with a coupling time of 0.1 ps. Pressure was kept at 1 bar using the Parrinello–Rahman barostat^35^ with a coupling time of 2.0 ps and an isothermal compressibility of 4.5 *×* 10*^−^*^5^ bar*^−^*^1^. Neighbor lists were updated every 10 steps using the Verlet scheme. Trajectories were saved every 100 ps, and, for steered or umbrella simulations, pull coordinates were saved every 10 ps.

### Weighted Histogram Analysis

Free-energy profiles along the reaction coordinate were reconstructed using WHAM, as implemented in the g_wham tool provided in GROMACS. Histograms were accumulated using the default binning scheme of g_wham, with window boundaries spanning *−*1.7 to +1.7 nm. WHAM convergence was achieved using a tolerance of 10*^−^*^6^.

Before the WHAM analysis, the sign of the reaction coordinate was assigned according to the position of Met1 relative to the reference plane defined above. The same sign transformation was applied consistently to both the sampled coordinate and the corresponding umbrella reference value. Thus, defining *d* = *σr* and *d*_0_ = *σr*_0_, with *σ* = *±*1 indicating the side of the plane, preserves the harmonic displacement exactly:

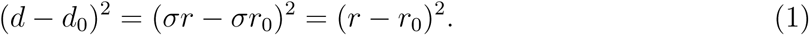

Frames that crossed the reference plane during a simulation were assigned to the corresponding signed branch. This procedure avoids artificial jumps in the signed coordinate while preserving, for every frame, the harmonic displacement and therefore the biasing potential applied during sampling. The unequal numbers of points in the histograms labeled *−*2 and +2 Å therefore reflect this branch classification rather than differences in the umbrella potentials used during the simulations.

The reconstructed PMFs were corrected for the radial Jacobian associated with using a three-dimensional distance as the reaction coordinate, according to^36^

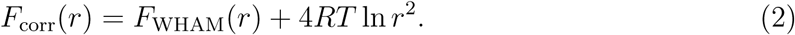

Equivalently, for the signed coordinate *d* = *σr*, the correction depends on the magnitude of the distance, *r* = *|d|*. This correction removes the geometrical contribution arising from the increase in configurational volume with radial distance.

Statistical uncertainties in the potential of mean force (PMF) were estimated using a jackknife resampling approach across replicates. The PMF was recomputed five times, each time omitting one of the five replicates for a given umbrella window (leave-one-out protocol). Only the production portions of each trajectory were included in the WHAM analysis.

### Structural Analyses

Standard structural descriptors were computed for all systems, including C*_α_* root-mean-square deviation (RMSD), radius of gyration (*R*_g_), and secondary structure content using the DSSP algorithm. ^37^ For holo systems, protein/ubiquitin interface areas were calculated using SASA data.^29,38^ To assess changes in surface chemistry upon unknotting, SASA values were weighted by the Wimley-White whole-residue hydrophobicity scale. ^39^ Binding free energies were estimated using MM-PBSA calculations performed with pyBindE^40,41^ (https://github.com/mms-fcul/PyBindE). Knot detection and classification were per-formed on every simulation frame using the Kymoknot software.^42^

## Results and discussion

### Simulation Convergence and Structural Stability

Before analyzing the sampling behavior along the reaction coordinate, we first assessed whether UCH-L1 remained structurally stable within each umbrella window. For all 17 umbrella windows (eight along the negative branch, eight along the positive branch, and the central window 00), we monitored several structural descriptors over time: C*_α_* RMSD, radius of gyration, secondary-structure content (DSSP), the number of helical residues within Helix 1, the number of threaded N-terminus residues, and the Cys90–His161 catalytic distance (Figures S1–S7 of the Supporting Information).

Across all windows, the RMSD remained within narrow fluctuation ranges and showed no evidence of unfolding or large-scale structural drift (Figure S1 of the Supporting Information). The radius of gyration was likewise stable (Figure S2 of the Supporting Information), indicating that the harmonic restraints did not significantly affect the UCH-L1 structure. DSSP analysis confirmed that although the global secondary-structure content was preserved throughout (Figure S3 of the Supporting Information), the number of helical residues in helix 1 was stable only in the umbrellas where the knot was still formed (Figure S4 of the Supporting Information). This is very clear from the number of threaded residues (Figures S5 and S6 of the Supporting Information), which are only well converged in knotted “positive” umbrellas. The Cys90–His161 distance is very difficult to equilibrate in only 100 ns per replicate (Figure S7 of the Supporting Information). Notwithstanding, we observe convergence, particularly after the initial 20 ns of the simulations, which were discarded.

### Characterizing the UCH-L1 Unknotting Process

During the unknotting process, UCH-L1 tends to continuously decrease the number of threaded residues at the N-terminus until the 5_2_ knot is fully disrupted, with the process completed in Umbrella *−*10 (Figure 2A). Although the process has little impact on the overall structure and stability of the protein, we observe some loss of secondary structure in Helix 1 (residues 10–21), in particular, when the knot is abolished (Figure 2B). This results from increased conformational fluctuations in the N-terminal segment, inducing additional strain at the start of the helical segment. This distortion does not occur when the protein is knotted. The positioning of the catalytic residues (distance between Cys90 and His161) is also easily affected by the integrity of the N-terminus and binding of ubiquitin.^1^ Hence, we applied a similar method to define a conformation as active (distance *<*5 Å; Figure S8 of the Supporting Information) and observed a trend in which UCH-L1 becomes less active upon unknotting (Figure 2C). After the Umbrella 00, we observe a transient alignment of the catalytic residues, although we have insufficient sampling of the N-ter segment position that drives this phenomenon.

**Figure 2:**
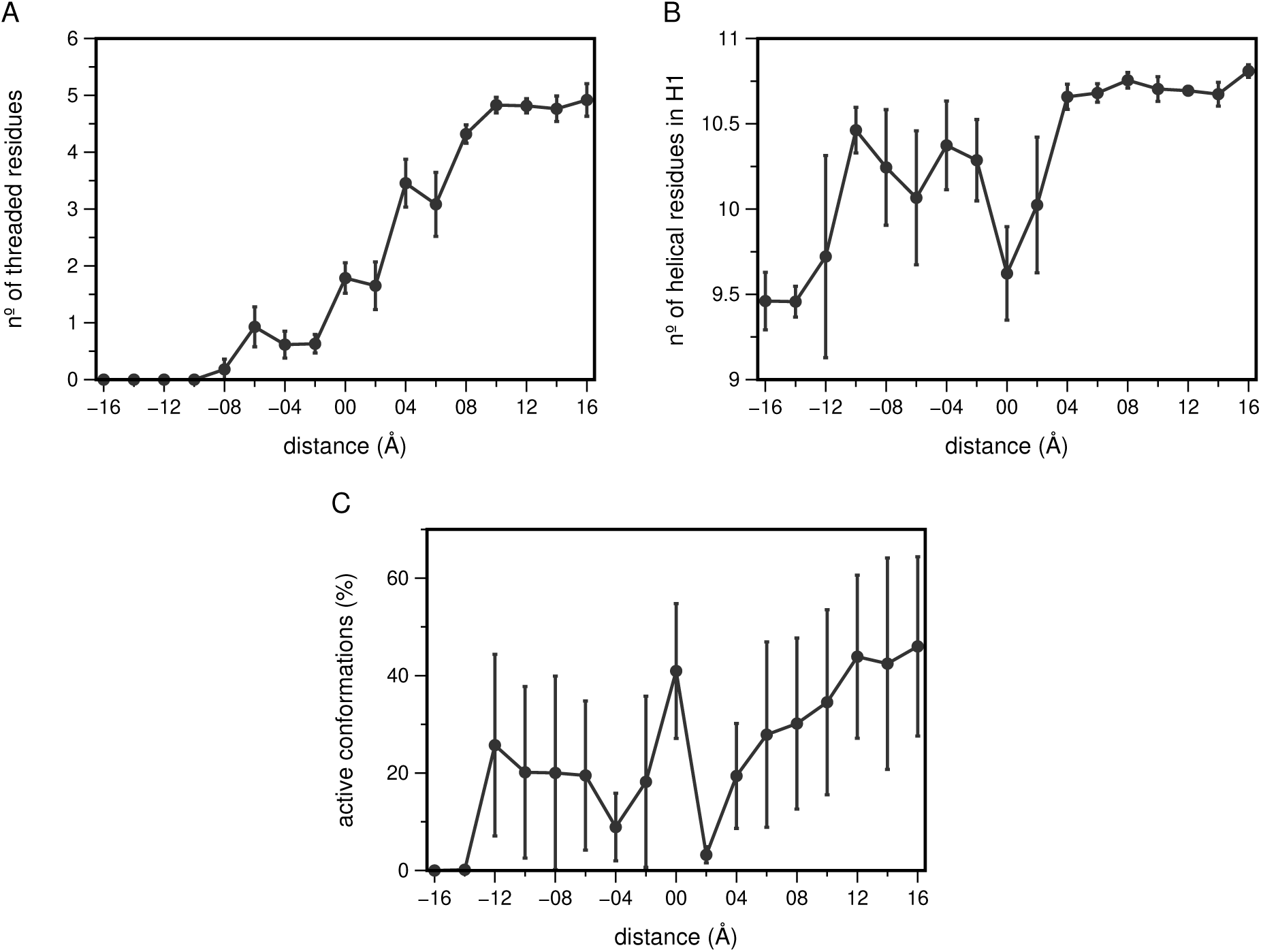
The number of threaded N-terminus residues (A), the number of helical residues in Helix 1 (B), and the percentage of active conformations (C), for each umbrella sampling window. The negative distances correspond to distances in the unknotted region of the collective variable. Active conformations correspond to distances *<*5 Å between Cys90 and His161 (Figure S8 of the Supporting Information).

We also monitored sampling along the reaction coordinate to ensure that neighboring US windows showed sufficient overlap for accurate WHAM reweighting calculations (Figure 3A). All 17 windows display distance histograms centered at the intended target values with substantial overlap across the full reaction-coordinate range (from *−*16 Å to +16 Å). The central region (Umbrellas *−*2, 0, and +2) shows modestly broader distributions relative to peripheral windows, reflecting the increased structural heterogeneity characteristic of conformations near the knotting/unknotting transition and the ability of the N-ter (reference) group to jump between the positive and negative regions of the collective variable. This was expected and indicates that these windows sample multiple relevant regions. However, it also poses a challenge in correctly assigning each conformation to the appropriate umbrella. When the N-terminus of UCH-L1 crosses under the reference loop, it jumps, for instance, between the *−*2 and +2 Umbrellas or vice versa. This needs to be corrected before the reweighting step to avoid misclassification and incorrect energy penalties. We accounted for these special cases by assigning a positive (+) or negative (-) value based on the position relative to a reference plane that intersects the gate loop’s center of mass.

**Figure 3:**
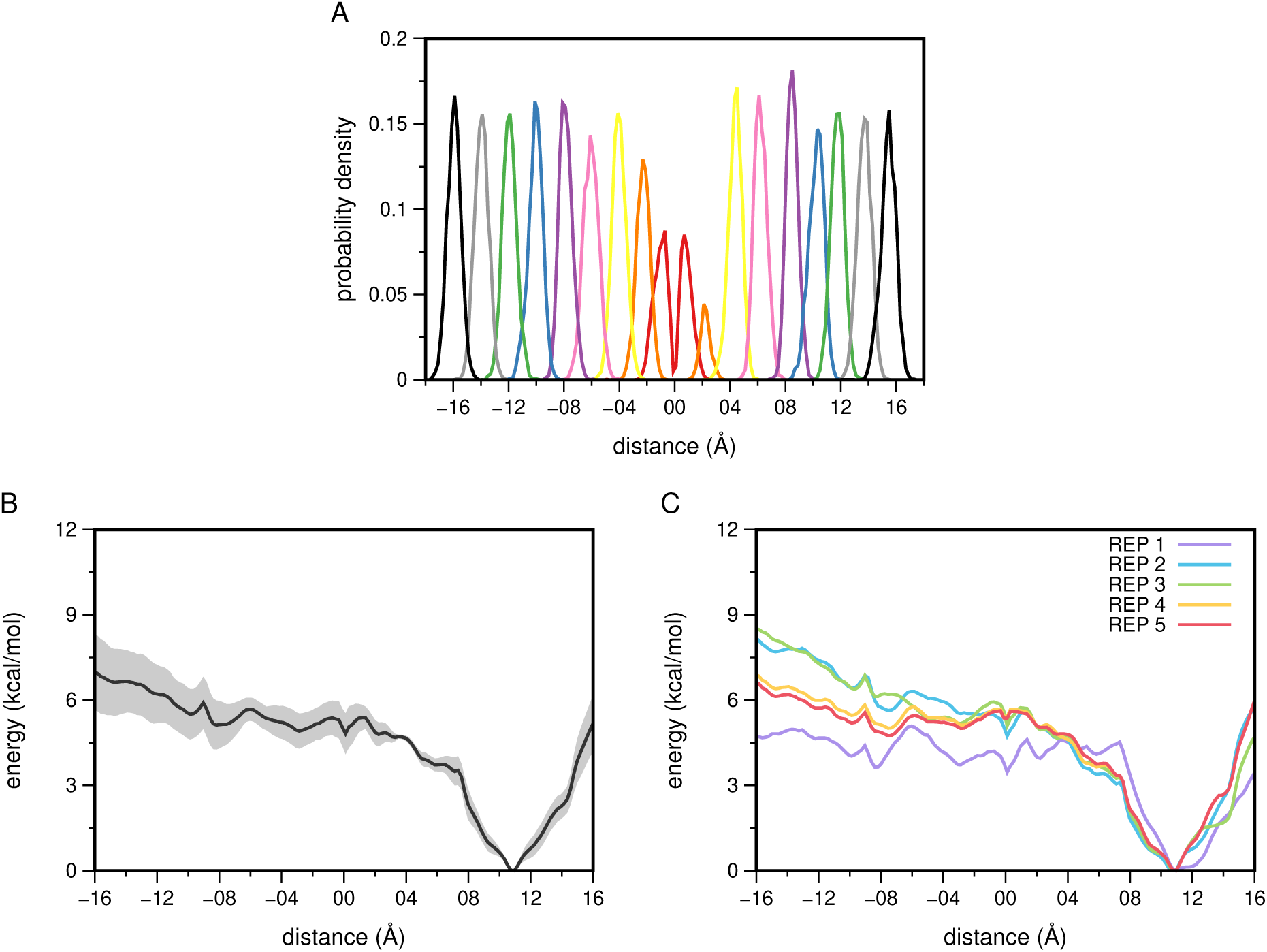
Distance histograms for all US windows (A) and the potential of mean force (PMF) for UCH-L1 unknotting, global (B) and per replicate (C). The gray-shaded region in the global PMF represents the standard error of the mean from the replicate data, estimated using the jackknife method. The negative distances correspond to distances in the unknotted region of the collective variable.

The free-energy profile along the constrained unthreading pathway sampled for UCH-L1 was estimated from the corrected potential of mean force (PMF) shown in Figure 3B. The native knotted region, centered near +11 Å, corresponds to the global free-energy minimum along this pathway. As the N-terminus moves towards and across the gate, the free energy increases, and the sampled unknotted conformations on the negative side of the coordinate remain approximately 5–7 kcal/mol above the native minimum. The profile does not exhibit a well-defined maximum at the gate that can be interpreted as the activation free-energy barrier or transition state for spontaneous unknotting. Instead, it indicates a thermodynamic preference for the native knotted topology along the constrained pathway sampled here.

The corrected PMF remains more irregular on the unknotted side of the gate, and the profiles become more replicate-dependent in this region (Figure 3C). This variability suggests that, after crossing the gate, the N-terminus explores a broader and more heterogeneous ensemble of conformations that is more difficult to sample reproducibly. Accordingly, the PMF should be interpreted as a pathway-dependent free-energy profile rather than an exhaustive description of the multiple unthreading geometries that contribute to the complete free-energy landscape. The pathway observed in our work is qualitatively consistent with the mechanism reported by Fonseka *et al.*, in which spontaneous unknotting in unconstrained atomistic simulations occurs through N-terminal unthreading via a gate. ^43^

### Impact of Unknotting in UCH-L1 Structure and Function

While the PMF provides a quantitative description of the relative free energy along the constrained unthreading pathway sampled here, the use of restraints and the simulation timescale of the umbrella-sampling protocol limit its ability to reveal the longer-term structural consequences of knot removal. To investigate these effects, we performed long unbiased MD simulations (10 *×* 500 ns) of the fully unknotted protein in both the apo and ubiquitin-bound (holo) states, examining how knot removal affects the stability, flexibility, and catalytic architecture of UCH-L1.

Analysis of RMSD, radius of gyration, secondary structure content, and the distance between Cys90 and His161 shows that both apo and holo unknotted variants retain overall fold stability across sub-microsecond simulations (Figure S9 of the Supporting Information). We observe very good convergence of all considered properties after the initial 200 ns, which were discarded. From the RMSF analysis, we observe that knot removal slightly increases the local flexibility of the UCH-L1 N-terminus compared with other regions of the protein (Figure 4). This is probably clearer in the holo form, where the impact is confined to the first *∼* 15 residues of the protein, with almost no contribution to global destabilization. These enhanced fluctuations in the apo form are consistent with the view that ubiquitin binding stabilizes UCH-L1. These results are also consistent with our previous ΔN truncation study,^1^ in which even minimal N-terminal deletions destabilized local contacts without inducing global structural rearrangements in the protein.

**Figure 4:**
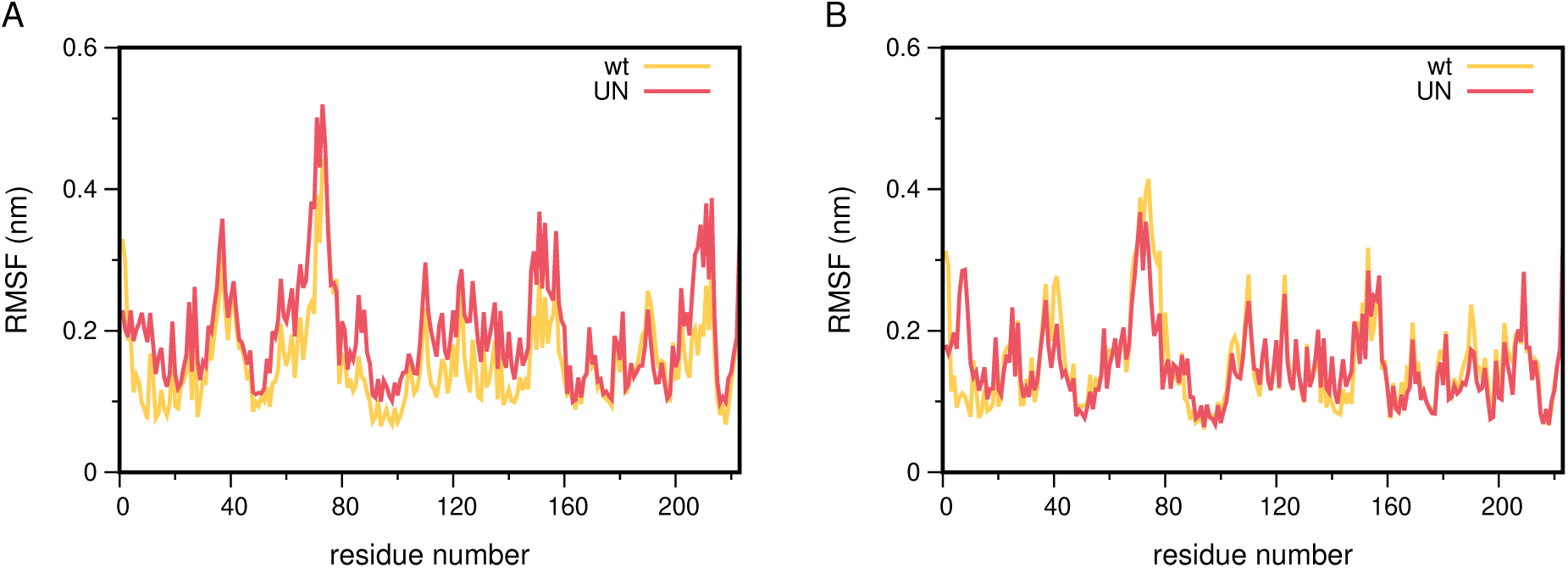
Root mean square fluctuations (RMSF) values of the apo (A) and holo (B) systems. The wild-type and unknotted forms are shown in yellow and red, respectively. Although ubiquitin is present in the holo systems, we present data only for UCH-L1 (223 a.a.).

The UCH-L1 knot anchors the N-terminus, and although ubiquitin binding partially compensates for its absence, the resulting structural distortions may still affect the protein’s active site. To examine how these perturbations could affect catalysis, we evaluated the distance between Cys90 and His161 (Figure S9D of the Supporting Information), which serves as a structural proxy for catalytic alignment. In our previous work, we observed that, for wild-type systems, catalytically competent geometries (*<*5 Å) were more frequently populated in the holo state than in the apo state (Figure 5). This trend is consistent with substrate-induced activation, whereby ligand binding brings together and stabilizes the reactive thiolate–imidazole dyad. ^1^ The unknotted variants exhibited similar profiles of active conformations to the wild type (Figure 5). Therefore, ligand binding was still able to restore the Cys90–His161 catalytic alignment after the knotted topology was removed. This effect is markedly different from what we observed for all truncated systems, some of which (ΔN2 and ΔN5) were confirmed experimentally, where even minor N-terminal deletions significantly reduced enzymatic activity.^1^

**Figure 5:**
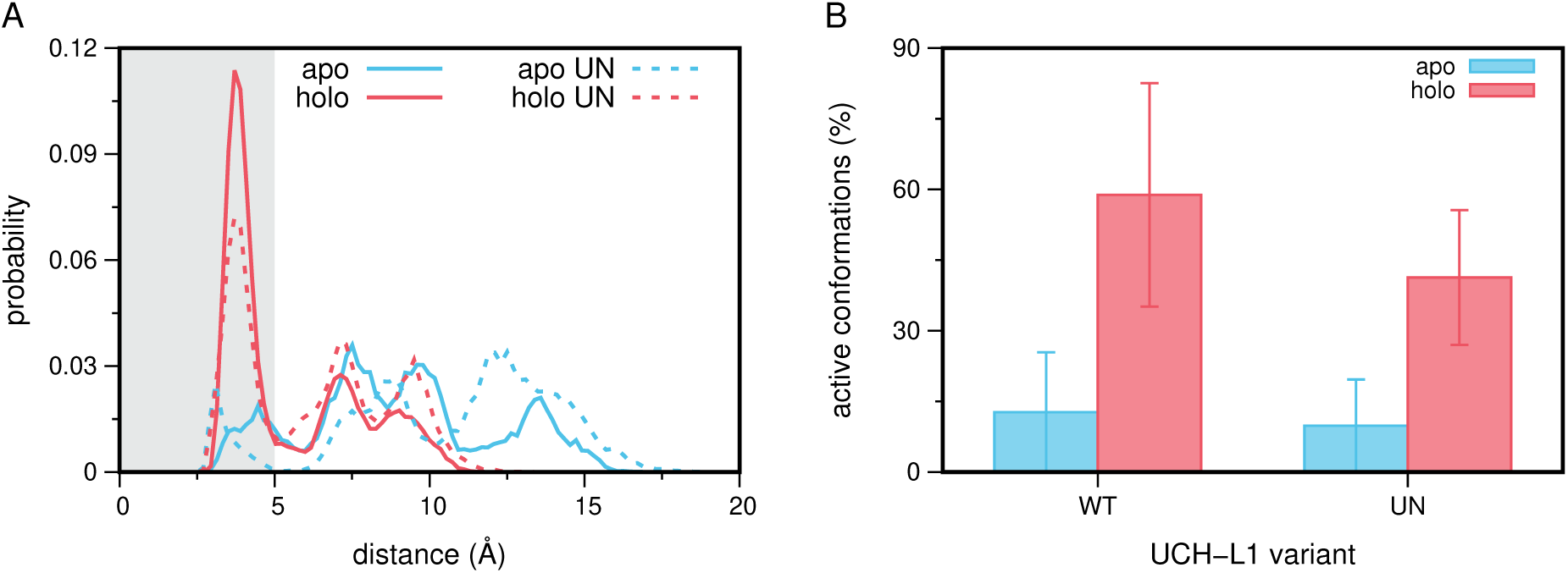
Probability distributions of the distance between Cys90 and His161 (A) and the percentage of active conformations (B) for all systems. In (A), the apo forms are shown in blue and the holo forms in red, with solid lines for the wild-type and dashed lines for the unknotted (UN) variant. The gray-shaded region indicates active conformations, defined as those within 5 Å. In (B), the apo and holo forms are shown in blue and red, respectively, for both the wild-type (WT) and unknotted (UN) variants.

Given that unknotting preserves the catalytic geometry of UCH-L1, we next examined whether it affects substrate binding. For this purpose, we performed MM-PBSA calculations to estimate ubiquitin binding free energies (Figure S10 and Table S1 of the Supporting Information). The unknotted variant exhibited significantly weaker binding than the wild type (*∼ −*62.1 *±* 1.4 kcal/mol vs *∼ −*75.5 *±* 1.8 kcal/mol), and although the high methodological uncertainty typically associated with MM-PBSA, our data support the interpretation that knot presence contributes to a higher substrate affinity, suggesting a possible topology-dependent feature.

To investigate how unknotting may alter ubiquitin binding to UCH-L1, we analyzed differences in solvent exposure between the wild-type and unknotted holo protein. Since a large variation in solvent exposure of one given residue may be more important than another, we used a metric called the hydrophobicity/hydrophilicity index, ^44^ which is obtained by multiplying the solvent-accessible surface area (SASA) by the Wimley–White whole residue hydrophobicity scale.^39^ The per-residue differences between the wild-type and unknotted forms can also be separated by hydrophilic (positive) and hydrophobic (negative) residues. Out of the 223 UCH-L1 residues (Figure S11 of the Supporting Information), only 11 show a significant change in solvent exposure upon unknotting (*|*ΔSASA*×*WW*| ≥* 0.5 Å^2^*·*kcal/mol) (Figure 6). Among hydrophobic residues, Ile8 is the only one that becomes significantly more exposed upon unknotting, consistent with its N-terminal position, which gains mobility after knot removal. RMSF analysis confirms this observation, showing that Ile8 transitions from a rigid, knotted conformation (RMSF of 0.10 nm in the wild-type) to a highly mobile state (RMSF up to 0.55 nm in the unknotted form) (Figure 4B). Additionally, this residue may also play an important role in ubiquitin binding, since unknotting leads to a significant decrease in the Δ*E*_VdW_ contribution to the MM/PBSA Δ*G*_bind_ energy (Table S1 of the Supporting Information). Among hydrophilic residues, several showed substantial changes in solvent exposure, with Glu7, Glu11, and Asp144 exhibiting the largest variations. In the wild-type, Glu7 is partially covered by the gate loop, while Asp144 is in close interaction with Lys4. Both residues lose their interaction partners upon unknotting and become more solvent-exposed, with substantially higher SASA*×*WW values in the unknotted form than in the wild type. In contrast, Glu11 exhibits a similarly large decrease in solvent exposure, becoming more buried upon unknotting. Notably, in the wild-type holo conformation, Glu11 is surface-exposed and the most flexible residue in the N-terminal region (RMSF of 0.16 nm). Upon unknotting, it becomes more rigid and buried because the mobile N-terminus collapses against the protein core, thereby restricting its solvent accessibility.

**Figure 6:**
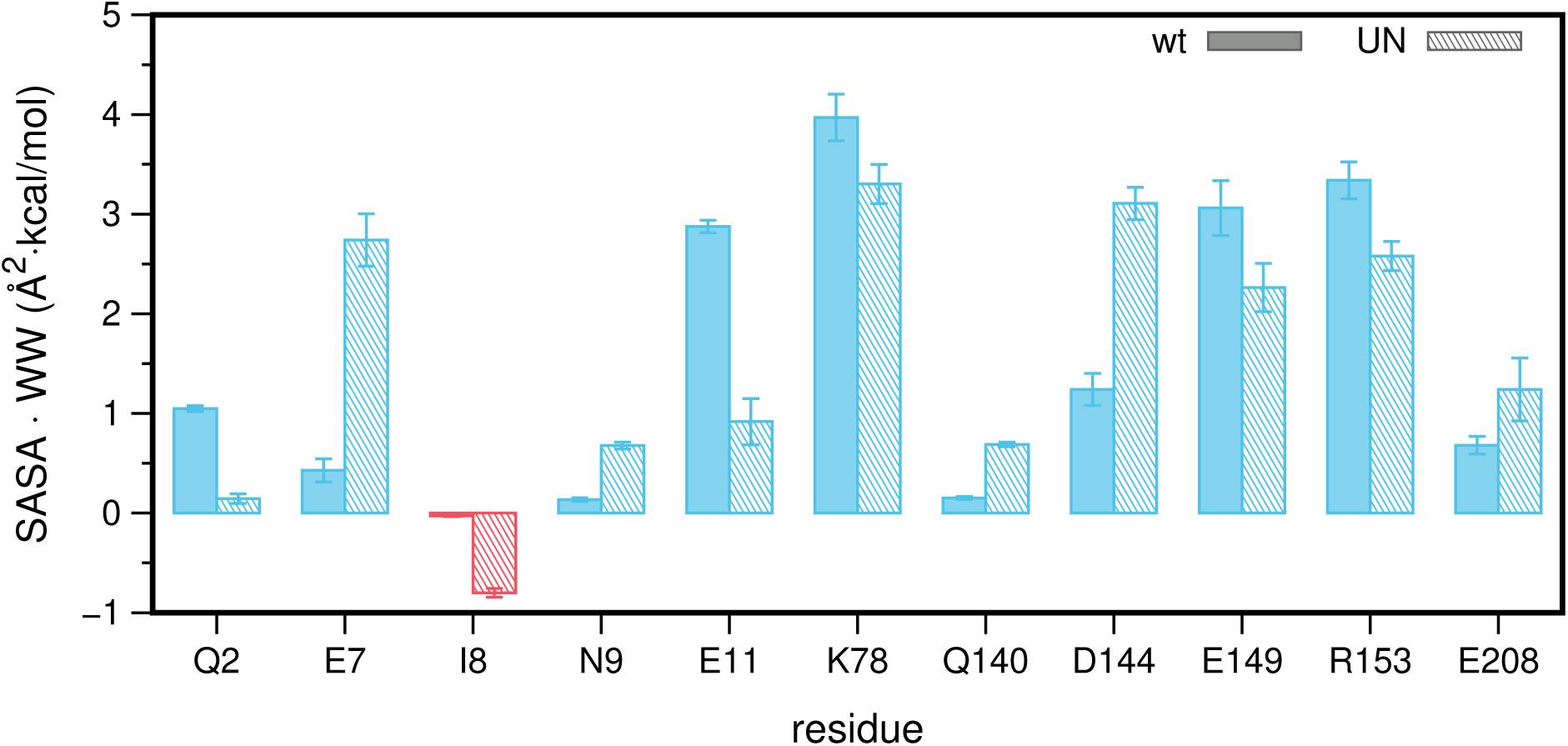
SASA*×*WW per residue for the 11 residues with significant differences between the wild-type and the unknotted UCH-L1 holo form (*|*ΔSASA*×*WW*| ≥* 0.5 Å^2^*·*kcal/mol). Hydrophilic residues (positive scale) are shown in blue and hydrophobic residues (negative scale) in red. Filled bars correspond to the wild-type (knotted) form and striped bars to the unknotted form. Error bars represent the standard error across ten replicates.

Together, these results show that the knotted topology in UCH-L1 does not dramatically alter global structural stability but imposes kinetic and structural constraints that seem to facilitate substrate binding. These results are consistent with our prior study, which showed that the structural integrity of the N-terminus is more important for enzymatic activity than the presence of the knot. ^1^ Nevertheless, the 5_2_ knot in UCH-L1 is not just a passive structural curiosity but also an apparent functional scaffold that could fine-tune substrate binding and contribute to the enzyme’s performance.

## Conclusions

This study quantifies the energetic and structural consequences of removing the 5_2_ knot from UCH-L1 while preserving its full-length amino-acid sequence. The calculated free-energy profile shows a clear thermodynamic preference for the native knotted topology along the constrained unthreading pathway sampled here, with the unknotted conformations lying approximately 5–7 kcal/mol above the native minimum.

Long unbiased MD simulations show that the unknotted variants retain the overall fold of UCH-L1, but display increased local flexibility, particularly at helix 1 in the apo state. Geometric analysis of the catalytic dyad further indicates that knot removal does not sub-stantially alter the sampling of catalytically competent conformations. MM-PBSA calculations predict less favorable ubiquitin-binding energetics upon unknotting, although both variants maintain stable ubiquitin-bound conformations throughout the simulations. The accompanying solvent-exposure analysis identifies hydrophobic and charged residues whose local environments change upon knot removal, and may contribute to the calculated difference between the two variants. Experimental measurements will be required to determine whether these computationally predicted changes translate into a measurable difference in ubiquitin-binding affinity.

Taken together, our previous truncation study^1^ and the present topology-preserving unknotting simulations suggest that the N-terminal segment and the associated knotted topology constitute a coupled structural-functional unit. Perturbations that disrupt either component alter functionally relevant properties, despite preserving the overall fold of the protein. These observations suggest that the native topology is not merely compatible with the architecture of UCH-L1, but may contribute to the local structural and dynamical framework underlying its interactions.

## Data and Software Availability

The GROMACS package is freely available software for MD simulations, available for down-load from https://manual.gromacs.org/documentation/2024.3/download.html. PyMOL v3.1 is also free software for molecular visualization and generating high-quality images. It can be downloaded from https://pymol.org. A zip file with all topologies, system configurations, and parameter files is also provided.

## Acknowledgement

We thank Sophie Jackson for fruitful discussions. We acknowledge financial support from Fundaçcão para a Cîencia e a Tecnologia through grant UI/BD/153055/2022 (BioSYS2 PhD Program) and projects 2023.15441.TENURE.006/CP00003/CT00004 and UID/04046/2025 (https://doi.org/10.54499/UID/04046/2025). This study was also supported by the European Union (TWIN2PIPSA - Twinning for Excellence in Biophysics of Protein Interactions and Self-Assembly, GA 101079147). Views and opinions expressed are those of the author(s) only and do not necessarily reflect those of the European Union or European Research Executive Agency (REA). Neither the European Union nor the granting authority can be held responsible for them.

## Supporting Information Available

Time series for RMSD, radius of gyration, secondary structure, number of threaded residues, and distance values (plus histograms) between Cys90 and His161 for both apo and holo systems for all windows in the umbrella sampling scheme. The RMSD, radius of gyration, secondary structure, distance between Cys90 and His161, MM/PBSA binding energies, and hydrophobicity indices from the long MD simulations of both end topologies (knotted and unknotted) in both the apo and holo states.

