## Supplementary Information File for "The Gordian Knot Enhances Ubiquitin Binding in UCH-L1"

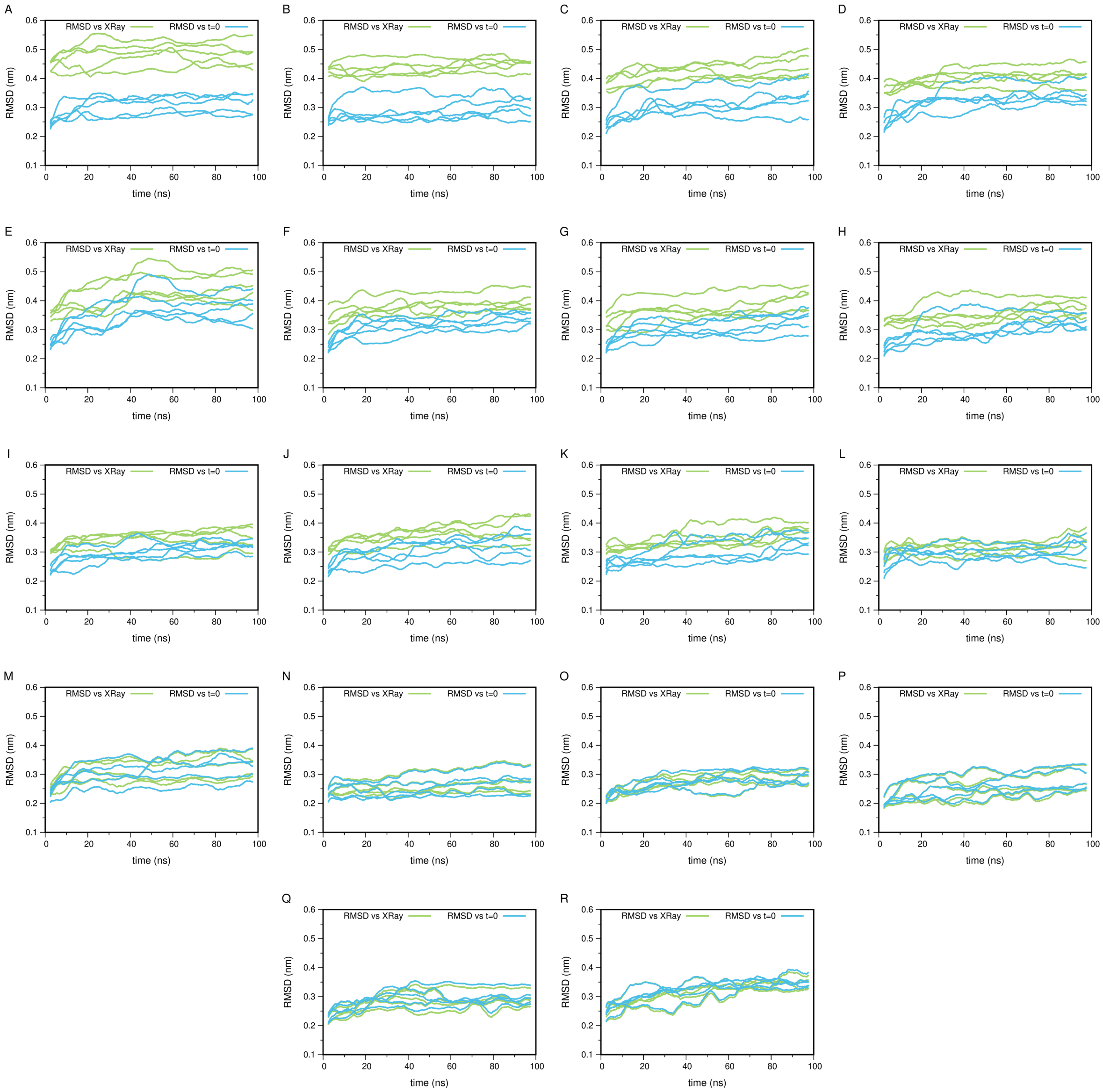


Figure S1: Root Mean Square Deviation (RMSD) across all umbrella sampling windows. Panels correspond to umbrella windows as follows: (A) −16 Å, (B) −14 Å, (C) −12 Å, (D) −10 Å, (E) −8 Å, (F) −6 Å, (G) −4 Å, (H) −2 Å, (I) 0 Å (unknotted), (J) 0 Å (knotted), (K) +2 Å, (L) +4 Å, (M) +6 Å, (N) +8 Å, (O) +10 Å, (P) +12 Å, (Q) +14 Å, (R) +16 Å. In blue, the UCH-L1's RMSD against the starting conformation, and in green, the RMSD against the crystallographic structure (PDB ID: 2ETL). The five replicates are shown. Data were averaged using a 5 ns floating window.

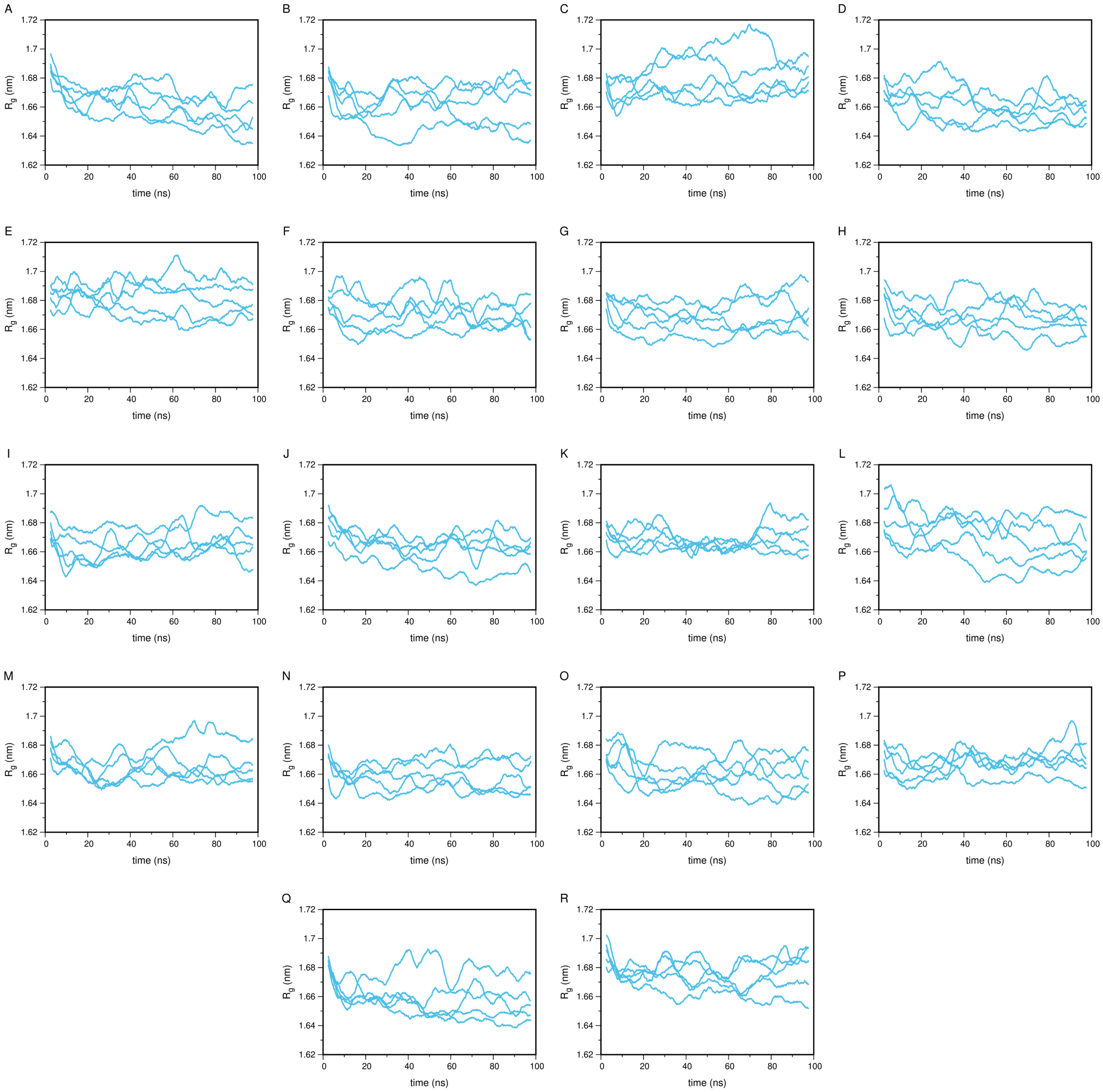


Figure S2: Radius of gyration across all umbrella sampling windows. Panels correspond to umbrella windows as follows: (A) −16 Å, (B) −14 Å, (C) −12 Å, (D) −10 Å, (E) −8 Å, (F) −6 Å, (G) −4 Å, (H) −2 Å, (I) 0 Å (unknotted), (J) 0 Å (knotted), (K) +2 Å, (L) +4 Å, (M) +6 Å, (N) +8 Å, (O) +10 Å, (P) +12 Å, (Q) +14 Å, (R) +16 Å. UCH-L1 is represented in blue. The five replicates are shown. Data were averaged using a 5 ns floating window.

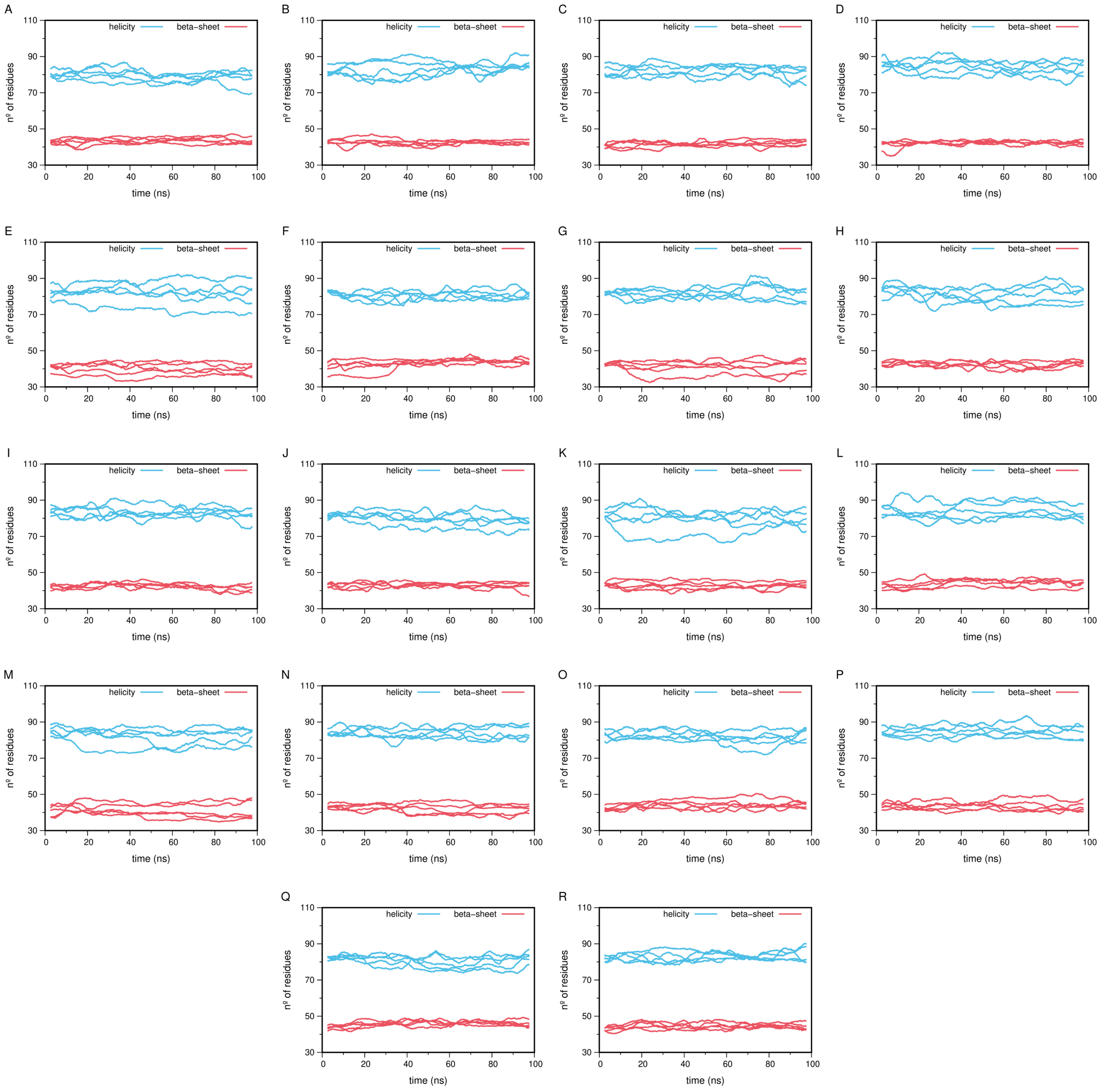


Figure S3: Secondary structure, using the DSSP criteria, across all umbrella sampling windows. Panels correspond to umbrella windows as follows: (a) −16 Å, (b) −14 Å, (c) −12 Å, (d) −10 Å, (e) −8 Å, (f) −6 Å, (g) −4 Å, (h) −2 Å, (i) 0 Å (unknotted), (j) 0 Å (knotted), (k) +2 Å, (l) +4 Å, (m) +6 Å, (n) +8 Å, (o) +10 Å, (p) +12 Å, (q) +14 Å, (r) +16 Å. $\alpha$-helices are represented in blue, and $\beta$-strands are represented in red. The five replicates are shown. Data were averaged using a 5 ns floating window.

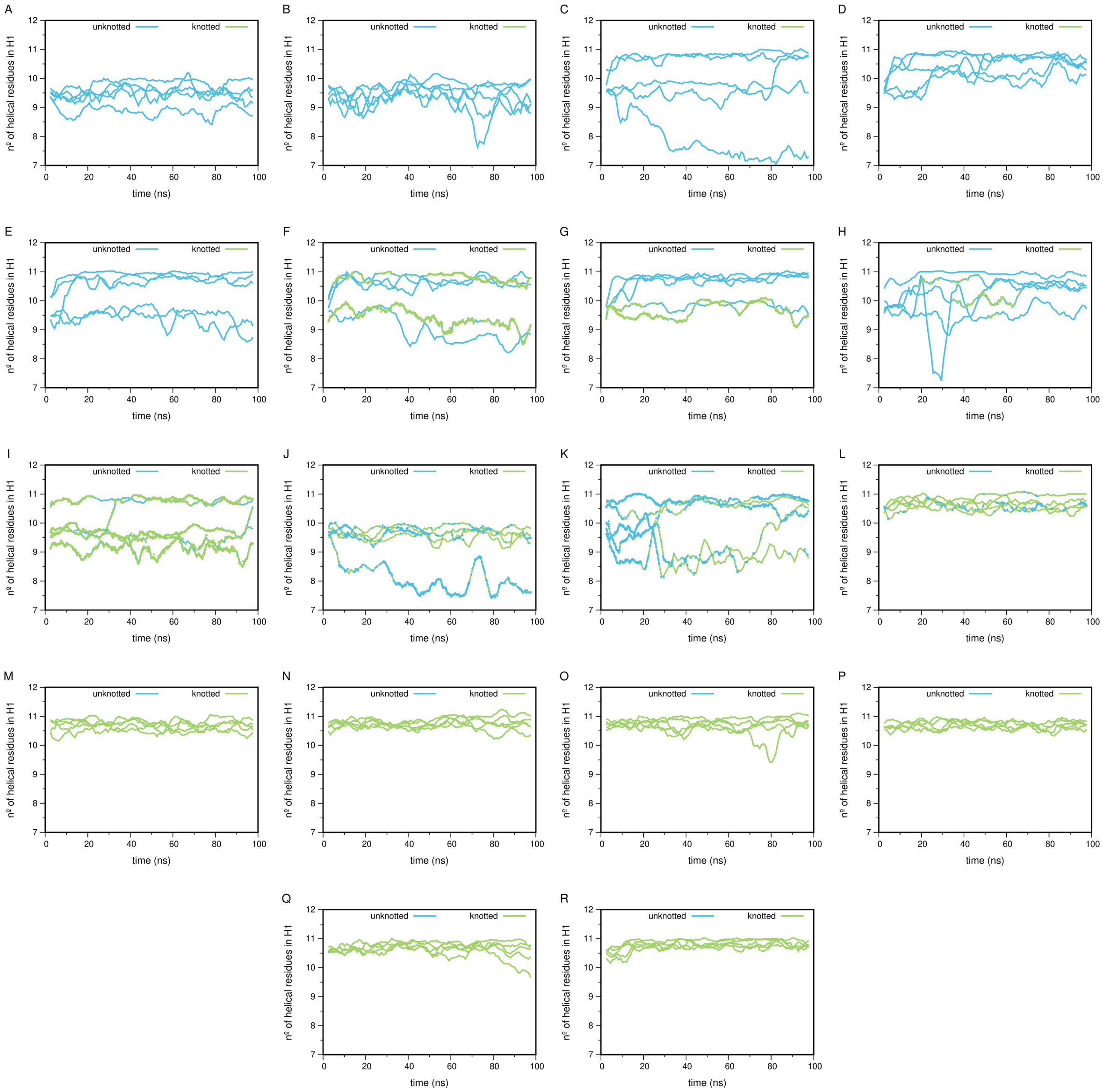


Figure S4: Number of helical residues present in helix 1 (H1), across all umbrella sampling windows. Panels correspond to umbrella windows as follows: (A) −16 Å, (B) −14 Å, (C) −12 Å, (D) −10 Å, (E) −8 Å, (F) −6 Å, (G) −4 Å, (H) −2 Å, (I) 0 Å (unknotted), (J) 0 Å (knotted), (K) +2 Å, (L) +4 Å, (M) +6 Å, (N) +8 Å, (O) +10 Å, (P) +12 Å, (Q) +14 Å, (R) +16 Å. UCH-L1 $\alpha$-helices are represented in blue for the unknotted topological state and in green for the knotted state. Because the reaction coordinate encodes only distance and not directionality, frames were classified as knotted or unknotted using a reference plane intersecting the gate center of mass. Color changes within a panel, particularly near the 0 Å window, reflect frames that crossed the gate during the simulation and were reassigned accordingly. The five replicates are shown. Data were averaged using a 5 ns floating window.

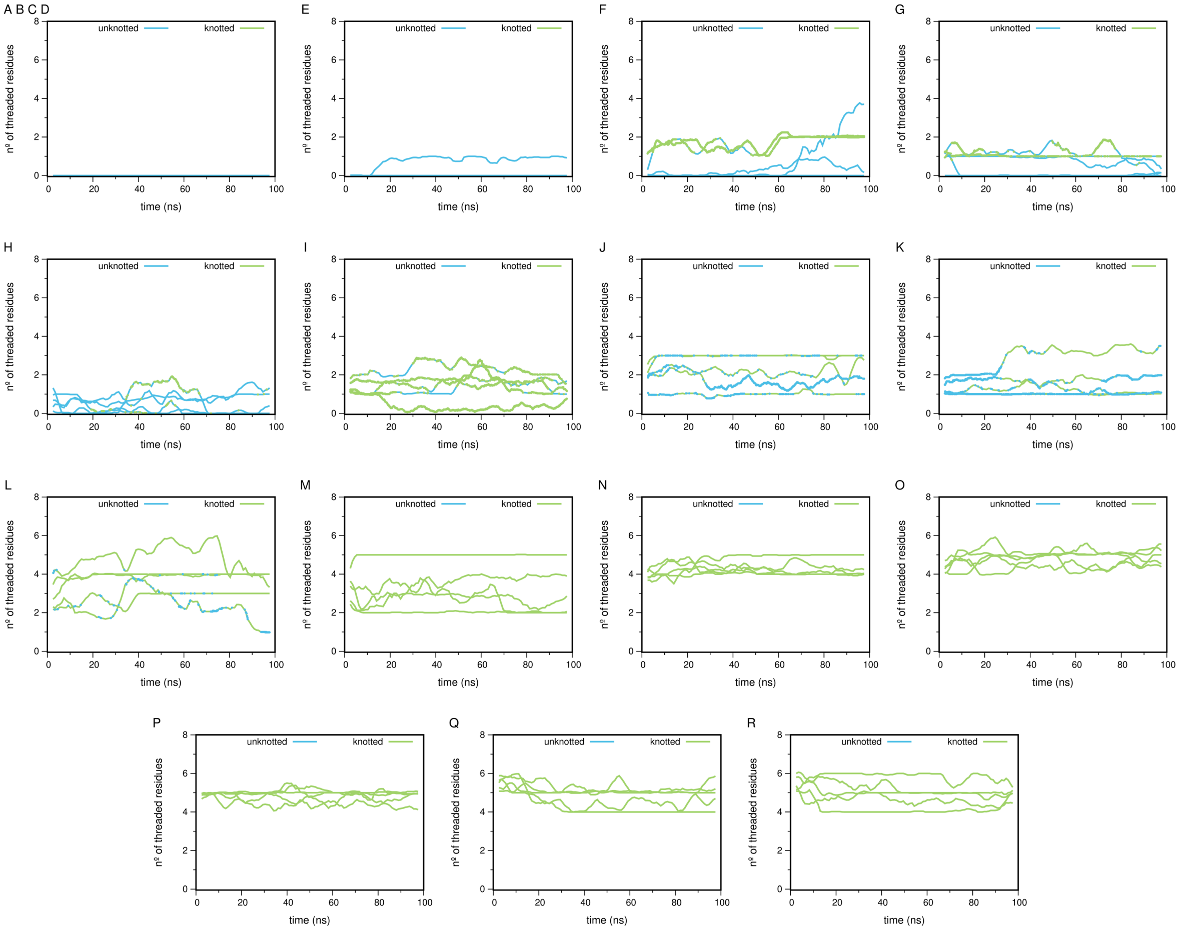


Figure S5: Number of threaded N-terminus residues across all umbrella sampling windows. Panels correspond to umbrella windows as follows: (A, B, C, D) −16 Å, −14 Å, −12 Å, and −10 Å(combined into a single panel as the number of threaded residues is consistently zero across all replicates in these windows, reflecting the fully unknotted state of UCH-L1), (E) −8 Å, (F) −6 Å, (G) −4 Å, (H) −2 Å, (I) 0 Å (unknotted), (J) 0 Å (knotted), (K) +2 Å, (L) +4 Å, (M) +6 Å, (N) +8 Å, (O) +10 Å, (P) +12 Å, (Q) +14 Å, (R) +16 Å. UCH-L1 is represented in blue for the unknotted topological state and in green for the knotted state. Because the reaction coordinate encodes only distance and not directionality, frames were classified as knotted or unknotted using a reference plane intersecting the gate center of mass. Some structural deformation can lead to knot assignment (Panel E), even when the energy penalty would make it highly unlikely (see Figure S6 for details). Color changes within a panel, particularly near the 0 Å window, reflect frames that crossed the gate during the simulation and were reassigned accordingly. The five replicates are shown. Data were averaged using a 5 ns floating window.

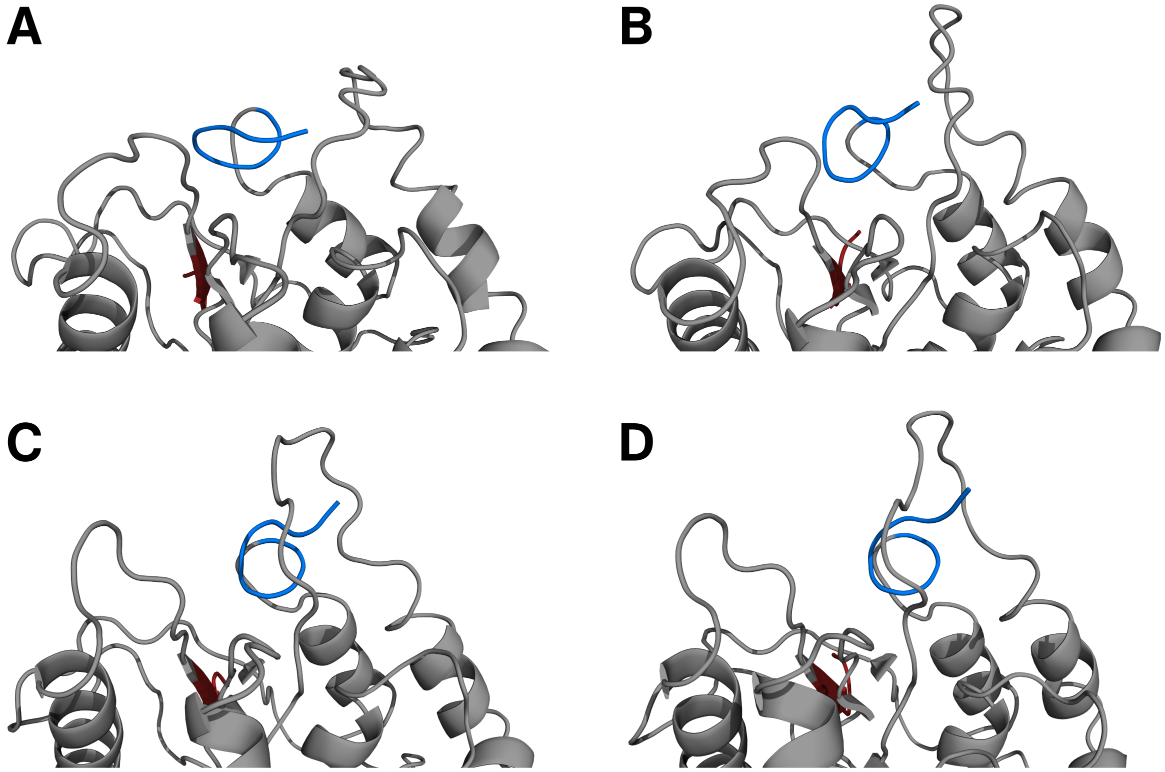


Figure S6: Illustration of the N-terminus position relative to the gate loop in umbrella window −8 (corresponding to an 8 Å unknotted displacement from the gate), for a representative replicate. Panels (A-D) show snapshots at $t$ = 0 ns, $t$ = 10 ns, $t$ = 30 ns, and $t$ = 70 ns, respectively. The snapshots show that the appearance of a single threaded residue in Figure S5E does not correspond to re-knotting of the protein, which would be energetically infeasible under the applied restraint. Instead, the gate loop opens sufficiently to allow the N-terminus to satisfy the imposed distance constraint of 8 Å from the center of mass of the gate residues (88, 140, 144, 147, 153, 155, and 157) while sitting slightly below the gate plane, and is therefore classified as having one threaded residue. The protein is represented as a gray cartoon. The N-terminal region (7 initial residues) is colored in blue, and the C-terminal region (7 final residues) is colored in red.

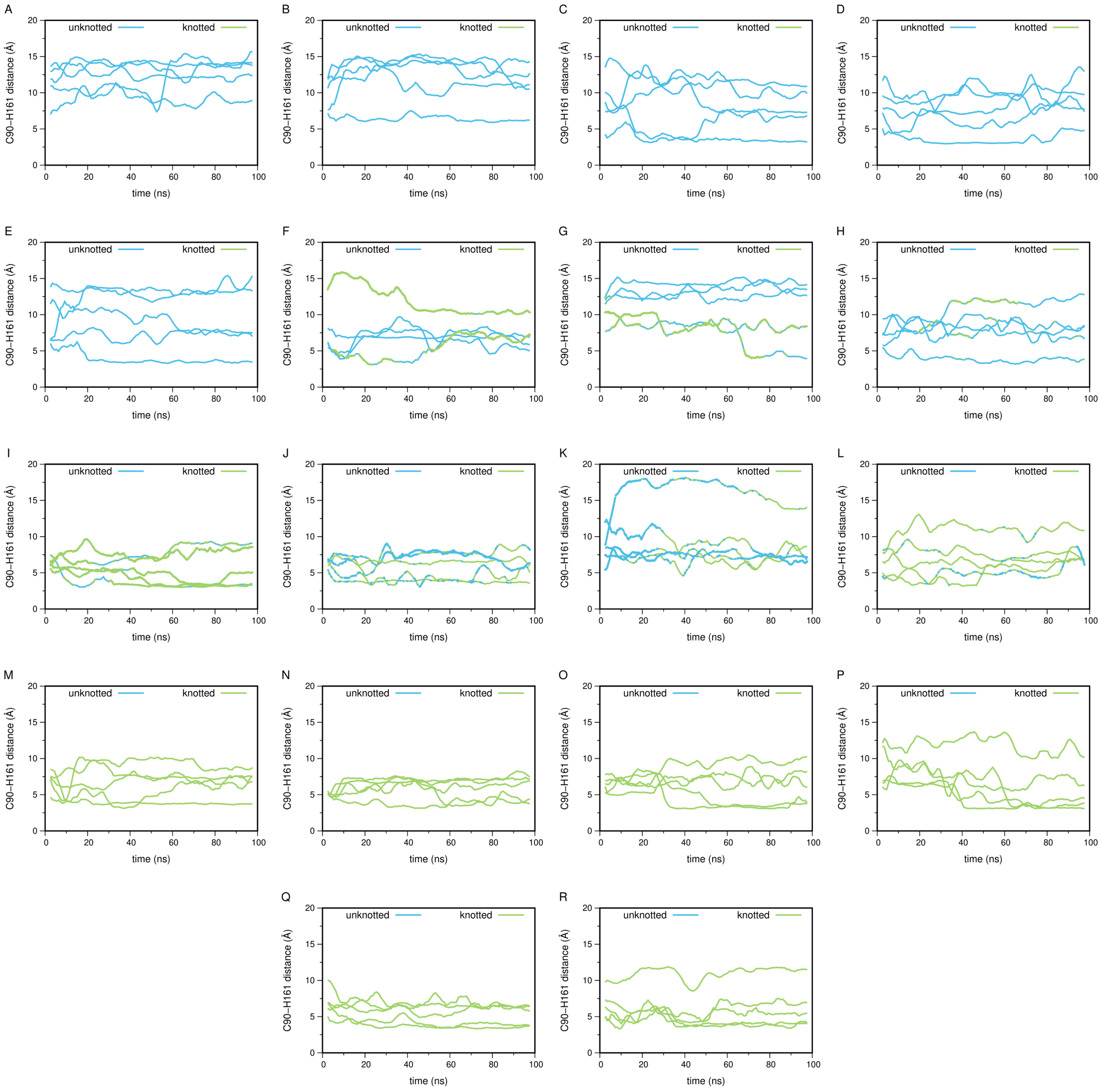


Figure S7: Distance between Cys90 and His161, across all umbrella sampling windows. Panels correspond to umbrella windows as follows: (A) −16 Å, (B) −14 Å, (C) −12 Å, (D) −10 Å, (E) −8 Å, (F) −6 Å, (G) −4 Å, (H) −2 Å, (I) 0 Å (unknotted), (J) 0 Å (knotted), (K) +2 Å, (L) +4 Å, (M) +6 Å, (N) +8 Å, (O) +10 Å, (P) +12 Å, (Q) +14 Å, (R) +16 Å. UCH-L1 is represented in blue for the unknotted topological state and in green for the knotted state. Because the reaction coordinate encodes only distance and not directionality, frames were classified as knotted or unknotted using a reference plane intersecting the gate center of mass. Color changes within a panel, particularly near the 0 Å window, reflect frames that crossed the gate during the simulation and were reassigned accordingly. The five replicates are shown. Data were averaged using a 5 ns floating window.

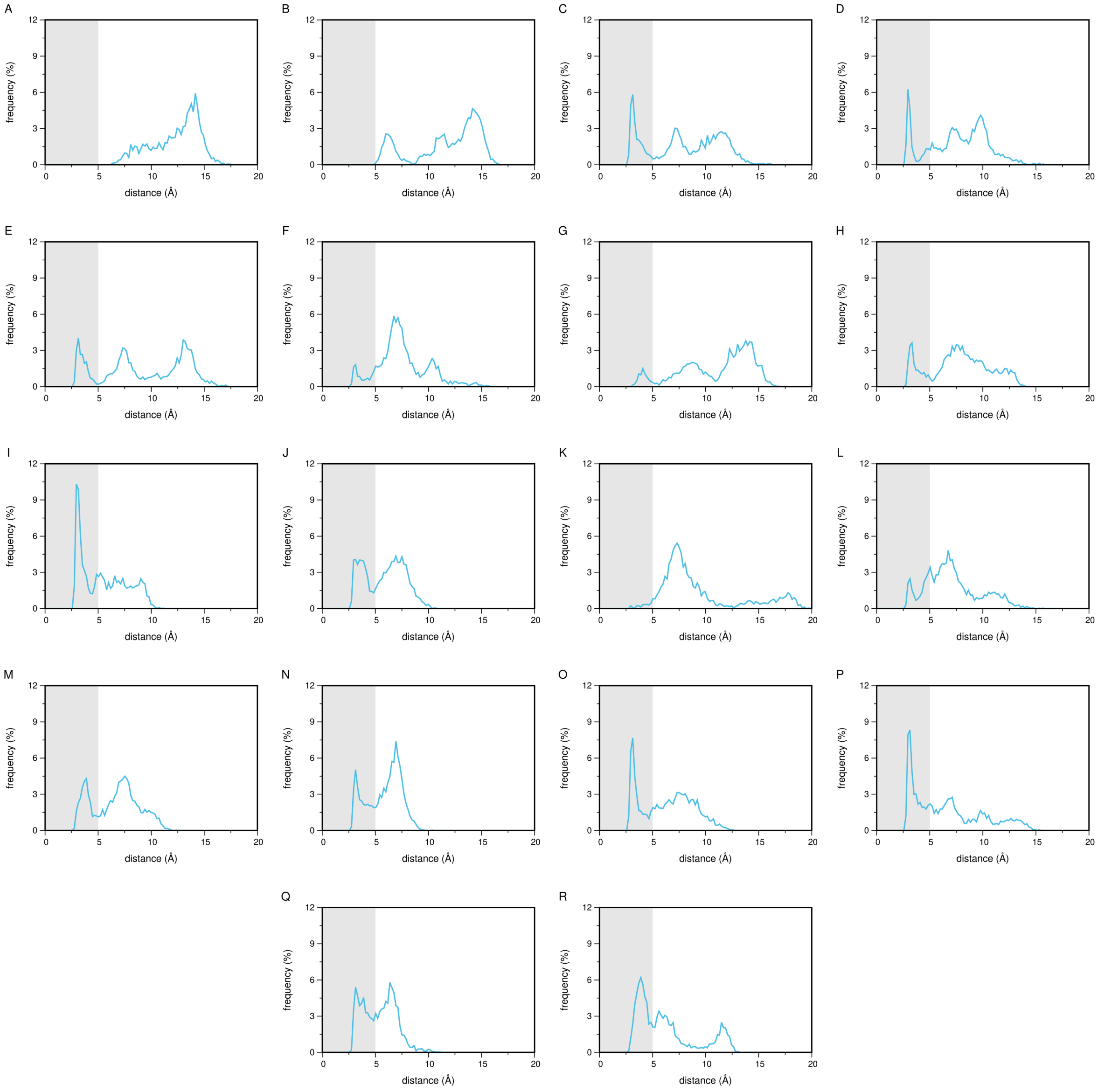


Figure S8: Distance histograms between Cys90 and His161, across all umbrella sampling windows. Panels correspond to umbrella windows as follows: (A) −16 Å, (B) −14 Å, (C) −12 Å, (D) −10 Å, (E) −8 Å, (F) −6 Å, (G) −4 Å, (H) −2 Å, (I) 0 Å (unknotted), (J) 0 Å (knotted), (K) +2 Å, (L) +4 Å, (M) +6 Å, (N) +8 Å, (O) +10 Å, (P) +12 Å, (Q) +14 Å, (R) +16 Å.

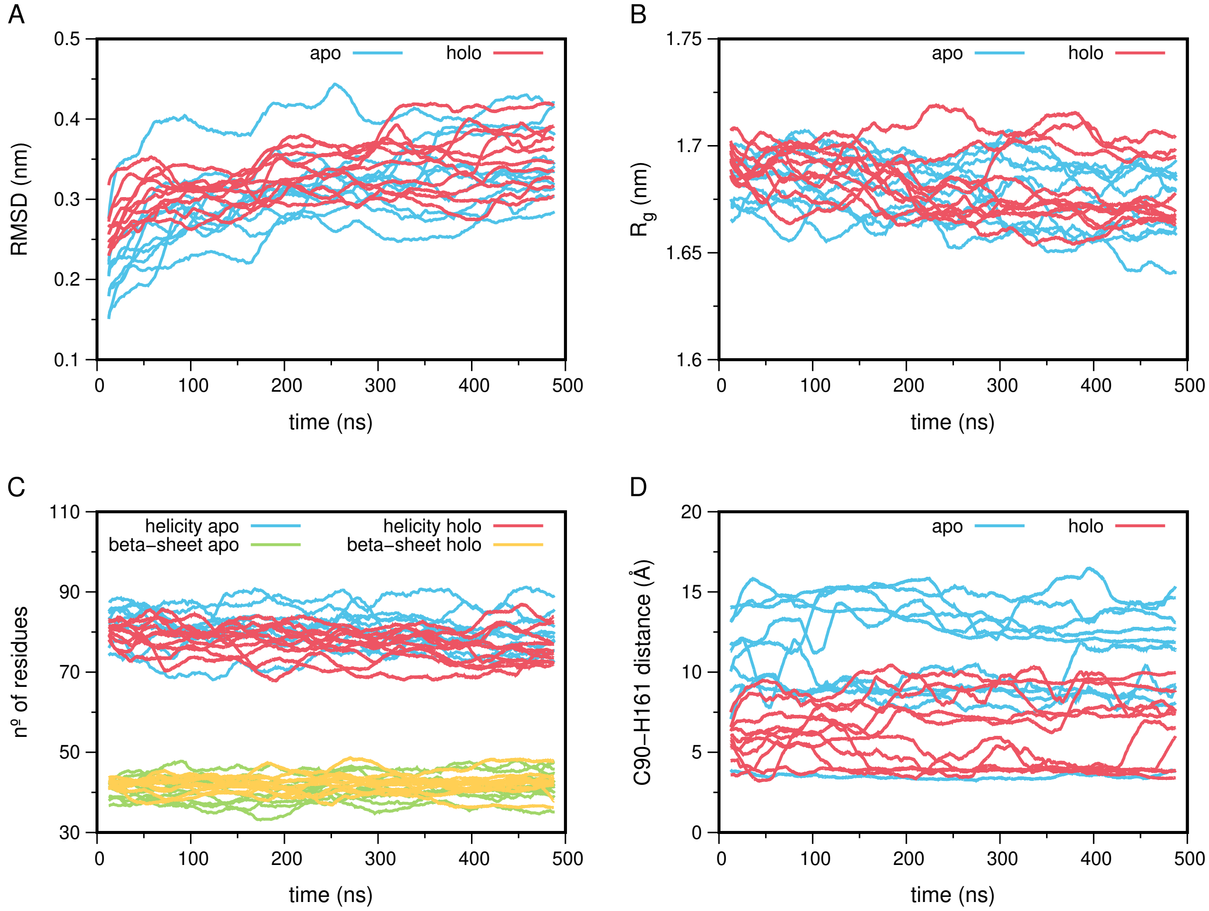


Figure S9: Root Mean Square Deviation (A), radius of gyration (B), secondary structure using the DSSP criteria (C), and the distance between Cys90 and His161 (D) for the wild-type UCH-L1. The apo state is shown in blue and the holo state in red. In (C), helices are shown in blue and red for the apo and holo states, respectively, and beta-sheets in green and yellow. Ten and five replicates are shown for the apo and holo states, respectively. Data were averaged using a 5 ns floating window.

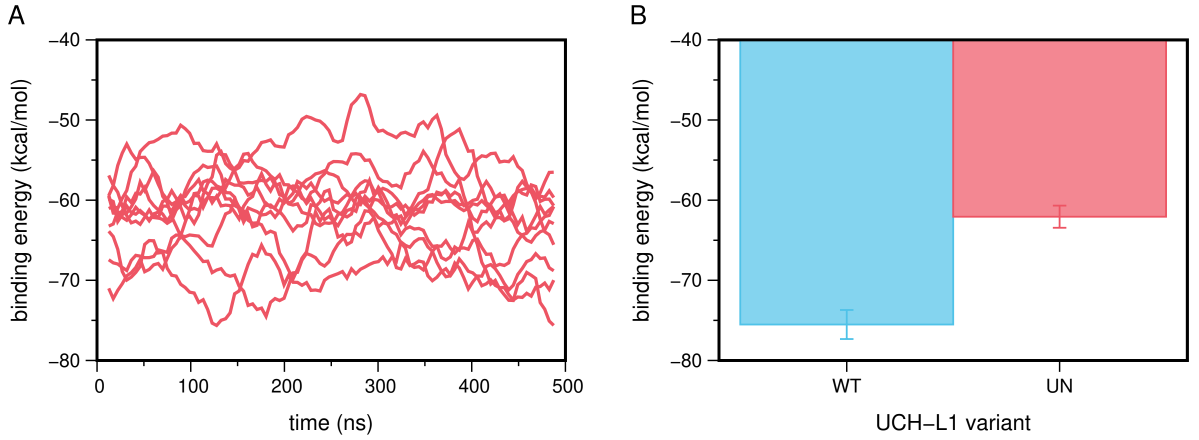


Figure S10: MM/PBSA binding free energy between UCH-L1 and ubiquitin. (A) Time evolution of the binding energy for the unknotted holo system, with ten replicates shown in red. Data were averaged using a 5 ns floating window. (B) Mean binding energy for the wild-type (blue) and unknotted (red) holo systems. Error bars represent the standard error across five and ten replicates for the wild-type and unknotted forms, respectively.

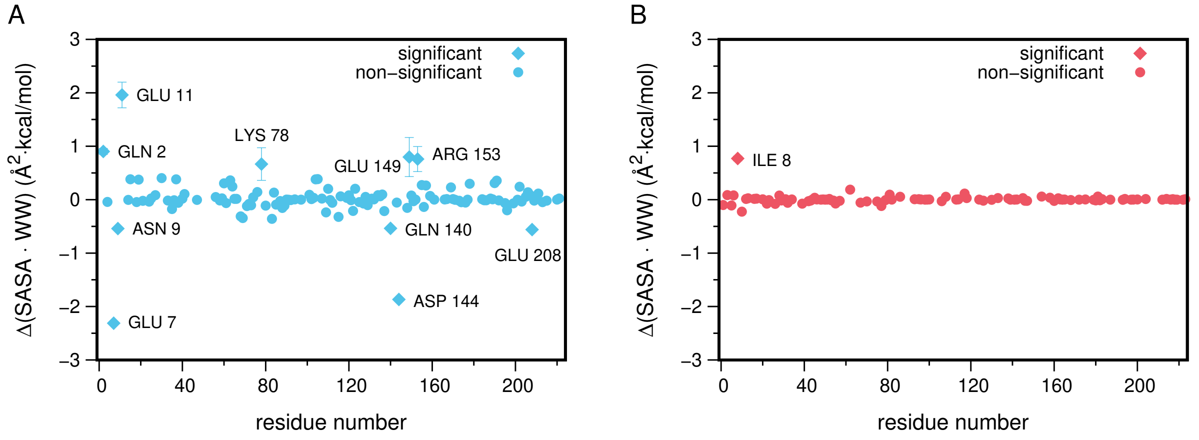


Figure S11: Difference in hydrophilicity (A) and hydrophobicity (B) between the wild-type and the unknotted UCH-L1 holo form. Positive/negative values indicate more exposed/buried residues upon unknotting. Residues with $|\Delta\mathrm{SASA}\times\mathrm{WW}| \geq 0.5$ Å$^2$·kcal/mol are considered significant (diamonds). Error bars represent the standard error across five and ten replicates for the wild-type and unknotted forms, respectively.

**Table S1: MM-PBSA energy contributions (kcal/mol) for the wild-type and unknotted UCH-L1 holo systems. Values represent the mean ± standard error.**

| Variant | $\Delta E_{\mathrm{VdW}}$ | $\Delta E_{\mathrm{Coul}}$ | $\Delta G_{\mathrm{nonpol}}$ | $\Delta G_{\mathrm{pol}}$ | $\Delta G_{\mathrm{bind}}$ |
|---|---|---|---|---|---|
| WT | $-105.1 \pm 3.9$ | $-139.1 \pm 7.7$ | $-12.9 \pm 0.4$ | $181.6 \pm 9.7$ | $-75.5 \pm 1.8$ |
| UN | $-82.9 \pm 2.5$ | $-103.0 \pm 8.2$ | $-11.1 \pm 0.4$ | $135.0 \pm 9.7$ | $-62.1 \pm 1.4$ |
